# *RRTF1* promotes touch-responses in *Arabidopsis* shoots independent of jasmonic acid

**DOI:** 10.64898/2026.03.08.710212

**Authors:** Sungkyu Park, Scott A. Finlayson, Chenxin Li

## Abstract

Plants acclimate to mechanical stimuli such as touch and wind via thigmomorphogenesis, a suite of developmental responses that alter their growth and architecture. However, the early signaling mechanisms translating mechanoperception into long-term morphological changes remain incompletely understood. We investigated the role of the rapidly touch-induced transcription factor RRTF1 (REDOX RESPONSIVE TRANSCRIPTION FACTOR 1) in these processes. Phenotypically, under aggressive mechanical stimulation, *rrtf1* mutant exhibited attenuated stunting (less height reduction). This suggests a key role for *RRTF1* in promoting thigmomorphogenic responses under severe mechanical stimuli, though the *rrtf1* mutant responded similarly to wild-type under gentle, repeated brushing. The alleviation of growth stunting in *rrtf1* was largely jasmonic acid (JA)-independent. Transcriptome analysis at 10 minutes post-touch revealed that *rrtf1* mutant maintained approximately 86% of wild-type touch-responsive gene expression. Nevertheless, *RRTF1* modulated specific regulons, partly through an interplay with WRKY transcription factors, as evidenced by altered TF binding motif enrichment in RRTF1-specific differentially expressed genes. We conclude that *RRTF1* acts as a modulator of early touch signaling in *Arabidopsis* shoots. It is not essential for the bulk of the initial transcriptional response but fine-tunes specific gene sets and plays a crucial role in calibrating long-term thigmomorphogenic development, particularly by promoting growth inhibition under severe mechanical stimulation. This study provides insights into the alleviation of touch-induced growth inhibition in *rrft1* mutant, which might be relevant to breeding for crops that are planted in high density and experience constant physical contact with neighboring plants.

## Introduction

Plants, in their natural settings, are continuously exposed to a diverse array of environmental stimuli, including both biotic and abiotic cues. Among these, mechanical stimulation−encompassing physical forces such as touch, wind, and rain−is a prevalent challenge to plants. Such stimuli trigger a series of adaptive mechanisms and biological responses that enable plants to cope with their environment and optimize fitness. Indeed, plants exhibit a wide range of developmental responses to mechanical stimulation (Braam and Davis, 1990; Braam, 2005). Periodic mechanical inputs can cause significant morphological changes, including reduced stem height and petiole length, altered stem mechanical properties, delayed flowering, and decreased leaf area (Jaffe, 1973; Chehab et al., 2012; Li et al., 2023; Zargar et al., 2022). This suite of responses to mechanical stimulation is termed thigmomorphogenesis (Jaffe, 1973). Thigmomorphogenesis has considerable impacts on important agronomic traits, often reducing plant height and grain yield while potentially increasing tiller number, shoot biomass, and stem biomechanical strength (Hindhaugh et al., 2021; Zargar et al., 2022). A clearer understanding of the mechanisms underlying thigmomorphogenesis could provide opportunities to fine-tune plant architecture and enhance crop productivity.

Thigmomorphogenesis can be viewed as an adaptive strategy enabling plants to mitigate varying levels of mechanical stress. This adaptation offers advantages, resulting in shorter, sturdier plants less prone to damage from environmental forces and physical handling, particularly in agricultural settings (Börnke and Rocksch, 2018), while ideally minimizing associated reductions in biomass and yield (Garner and Björkman, 1999; Graham and Wheeler, 2017). Additionally, mechanical stimulation can enhance plant disease resistance (Benikhlef et al., 2013; Matsumura et al., 2022). Given the slow development of thigmomorphogenic traits versus the rapid molecular responses to mechanical stimuli, a key challenge is to understand how prolonged and repeated stimulation translates into these lasting morphological changes. Therefore, it is important to induce robust and repeatable thigmomorphogenesis and to define the early signaling events that trigger touch responses. This knowledge may aid in developing strategies to mitigate any negative effects of thigmomorphogenesis on crop yield and to harness its benefits, such as increased stress resistance. An important step in this endeavor is the identification of key early signaling components.

The nature of plant responses to mechanical stimulation varies depending on the parameters of the stimulus, such as the intensity, frequency, and duration of the applied forces (Telewski and Pruyn, 1998; Bonnesoeur et al., 2016; Kašpar et al., 2017). Tissue specificity and the developmental stage at which stimulation occurs also play crucial roles in determining these responses (Biddington, 1986; Paul-Victor and Rowe, 2011). While investigations into how plants respond to mechanical forces have gained momentum at the molecular level, a complete understanding of the sophisticated molecular machinery for sensing mechanical cues remains a challenge. Although significant progress has been made in deciphering mechanical force signal transduction in *Arabidopsis* roots including receptor-like kinases and mechanosensitive ion channels, only a few signaling pathways relevant to shoot mechanical stimulation, such as jasmonic acid (JA)-dependent and -independent pathways, have been identified (Monshausen et al., 2009; Yamanaka et al., 2010; Chehab et al., 2012; Shih et al., 2014; Van Moerkercke et al., 2019; Fang et al., 2021; Mousavi et al., 2021; Darwish et al., 2022).

Mechanical stimulation responses comprise a cascade of complex signaling pathways beginning with the perception of signals in cellular structures such as cytoskeleton and actin filaments, and membrane-bound mechanosensitive ion channels (Telewski, 2006). Mechanical perturbations trigger immediate, transient, and dose-dependent elevations of cytosolic Ca²⁺, designating the ion as an early mechanosensory signal (Toriyama and Jaffe, 1972; Knight et al., 1991; Knight et al., 1992). These bursts activate downstream cascades involving reactive oxygen species and the hormones ethylene, auxin, and jasmonic acid (Biro and Jaffe, 1984; Monshausen et al., 2009; Monshausen and Haswell, 2013; Sassi and Traas, 2015; Van Moerkercke et al., 2019). The propagation of these intracellular Ca²⁺ waves sets off phosphorylation cascades, including MAPK cascades and widespread transcriptomic reprogramming (Wang et al., 2018; Matsumura et al., 2022). Extensive research has focused on the transcriptional reprogramming underlying touch responses (Chehab et al., 2012; Van Moerkercke et al., 2019; Darwish et al., 2022; Li et al., 2023). Despite significant progress in characterizing mechanosensing, the specific signaling components and pathways that translate mechanical signals into thigmomorphogenesis remain incompletely understood. Elucidating these early signaling events in aboveground plant parts is crucial for understanding how transient mechanical stimuli drive long-term morphological adaptation.

Mechanical stimulation involves a complex interplay of signaling molecules, including cytosolic Ca²⁺, ROS, nitric oxides (NO), and phytohormones (Garcês et al., 2001; Van Breusegem et al., 2001; Mori and Schroeder, 2004; Miller et al., 2009; Herbette et al., 2011; L’Haridon et al., 2011; Benikhlef et al., 2013; Fichman et al., 2019; Zandalinas et al., 2020). Among these, phytohormones are central to driving the core morphological and developmental changes associated with thigmomorphogenesis (Chehab et al., 2012; Börnke and Rocksch, 2018; Van Moerkercke et al., 2019; Xu et al., 2019; Darwish et al., 2022; Darwish et al., 2023; Li et al., 2023). While various hormones are implicated, thigmomorphogenesis has been proposed to result primarily from crosstalk between jasmonic acids (JA) and gibberellins (GA) (Chehab et al., 2012; Lange and Lange, 2015; Wang et al., 2023). For example, mechanical stimulation elicits a rapid increase in JA levels in both sorghum and *Arabidopsis* (Chehab et al., 2012; Liu and Finlayson, 2019; Van Moerkercke et al., 2019; Li et al., 2023; Park et al., 2024), implicating JA as an early touch signaling component.

AP2/ERF TFs, one of the largest transcription factor families in plants, play essential roles in growth, development, hormone regulation, and stress responses (Xie et al., 2019). Certain APETALA2/Ethylene Responsive Factor (AP2/ERF) transcription factor genes exhibited rapid and robust induction in response to mechanical stimulation in both sorghum buds and stems, alongside changes in phytohormone levels (Park et al., 2024). Consistently, in *Arabidopsis*, gentle touch stimuli trigger rapid transcriptomic responses, including the induction of several ERF genes within 25 minutes (Van Moerkercke et al., 2019; Darwish et al., 2022). Among the early-induced ERF genes, *RRTF1 (Redox Responsive Transcription Factor 1)* shows a particularly strong and rapid response to mechanical perturbations, with its expression typically peaking within an hour (Liu and Finlayson, 2019; Van Moerkercke et al., 2019; Darwish et al., 2022). Recent findings indicate that RRTF1 interacts with MED25, a subunit of the Mediator transcriptional co-activator complex and a key regulator in JA signaling, and JAZ9 repressor (Ou et al., 2011; Zhang et al., 2019; Zhai et al., 2020). While RRTF1 expression is induced by JA, the precise mechanism of JA in this induction is complex (Cai et al., 2014). Given the multifaceted roles of MYCs in JA pathways (Van Moerkercke et al., 2019; Chico et al., 2020; Liu et al., 2019), and the association of this JA-dependent MYC network with RRTF1, it is important to understand if and how RRTF1-mediated thigmomorphogenesis depends on JA signaling. RRTF1 homologs are found in various plants, suggesting a conserved role in mechanical signaling. In sorghum, a *RRTF1* homolog was highly induced in tiller buds after leaf removal (Liu and Finlayson, 2019; Park et al., 2024). Studies in other cereals like barley, oat, and wheat also show rapid *RRTF1* homolog induction by mechanical stimulation, albeit with some species-specific timing variations (Darwish et al., 2023). This points to a conserved role for RRTF1 in mediating mechanical responses across diverse plant species.

The current study generated, compiled, and analyzed transcriptome profiles to identify key regulators of plant mechanical responses. Through screening touch-induced transcriptomes across various time frames, we identified a key role for *RRTF1* in promoting thigmomorphogenic responses under severe mechanical stimuli. We also uncovered that RRTF1 and WRKY transcription factors share a regulon, which includes a subset of the core touch-responsive genes.

## Results

### ERF transcription factors, including RRTF1, are rapidly induced by mechanical stimulation

Touch-responsive genes are rapidly induced by mechanical stimulation, including water spray, wind, and brushing. However, reproducing consistent results across different experimental setups is challenging because mechanical responses vary with the frequency and magnitude of the applied forces. While various transcriptome profiles in response to mechanical stimuli have been reported, only a few studies have detailed early transcriptional changes (within 30 minutes) following mechanical stimulation. For instance, studies in *Arabidopsis* and sorghum have revealed dynamic and widespread changes in transcription profiles between 5 and 30 minutes post-mechanical stimulation (Van Moerkecke et al., 2019; Wang et al., 2023; Park et al., 2024). Given this extreme sensitivity of this response, capturing true early mechanical signals requires strict isolation from background noise.

To elucidate the genetic factors underlying mechanical responses and early transcriptional changes, RNA-seq analysis was conducted on *Arabidopsis* subjected to mechanical stimulation. 10- to 12-day-old *Arabidopsis* seedling shoots were harvested at 5, 10, and 30 minutes after being touched by a paintbrush. Seedlings were grown carefully and remained unstimulated until the touch treatment. Naive *Arabidopsis* shoots were gently touched four times with a soft paintbrush. Untouched plants were harvested as controls after the touched plants to limit any effects of the harvesting process on touch responses. To identify differentially expressed genes following this touch treatment, cleaned reads (approximately 99% of the 10-15 million reads obtained) were mapped to the *Arabidopsis* reference genome (TAIR10, Lamesch et al., 2012). The number of reads, mapped reads, and the percentage of mapped reads in each replicate are shown in Supporting Information Table S1. In total, 20,201 genes were identified as expressed, based on the criterion of having raw counts > 10 in at least three samples. Principal component analysis (PCA) plot of the read counts transformed by variance stabilizing transformation (VST) showed that biological replicates clustered together within each time point. The PCA plot also revealed a clear separation between time points along the PC1 and PC2 axes following touch (Fig. 1a).

**Fig. 1.**
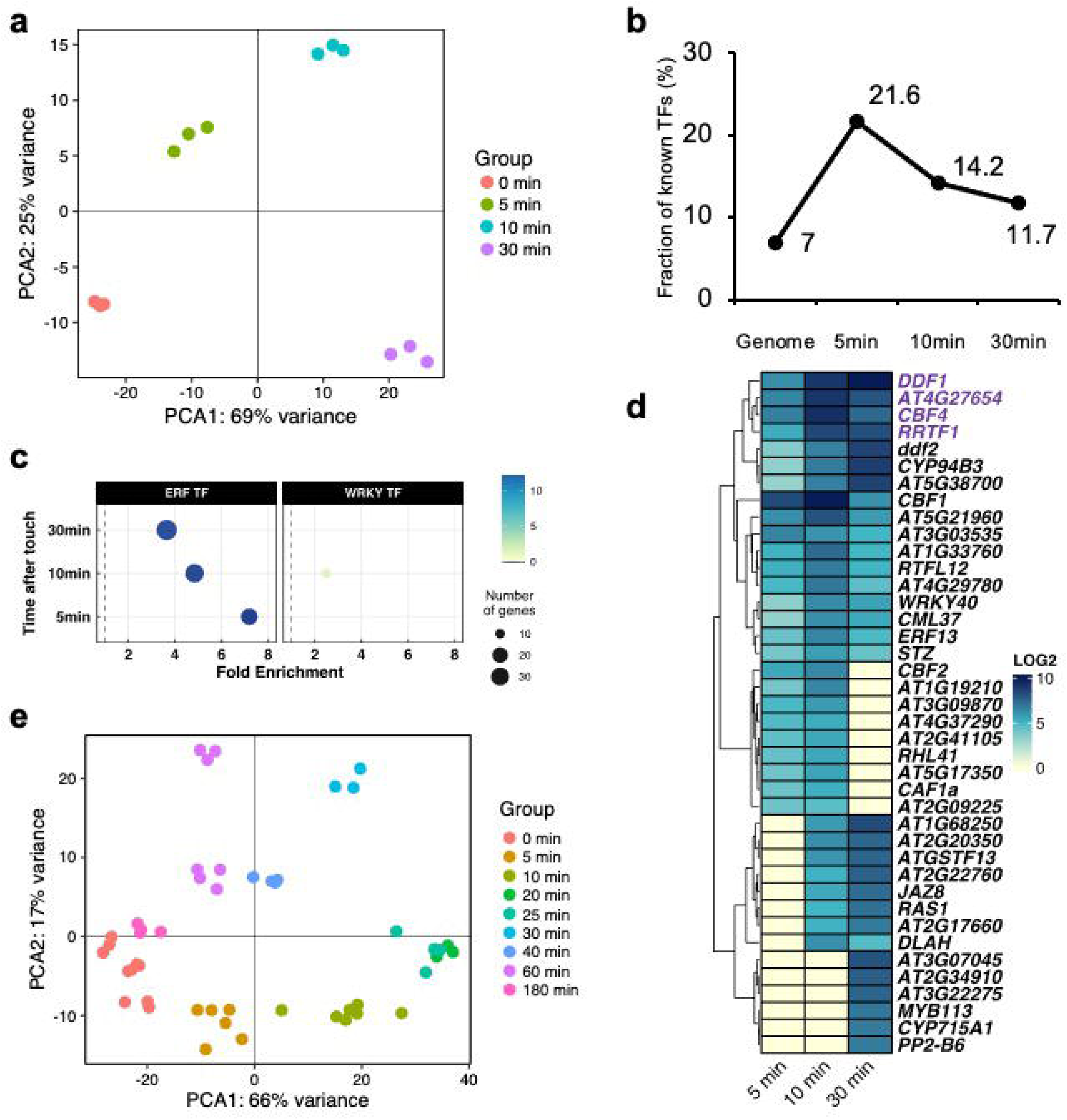
Rapid induction of ERF transcription factors, including *RRTF1,* in response to mechanical stimulation, (a) Principal component analysis (PCA) of differentially expressed genes (DEGs) at 5, 10, and 30 min after touch, (b) Proportion of DEGs encoding known transcription factors (TFs) at 5, 10, and 30 min after touch, (c) TF enrichment analysis of touch-responsive DEGs. (d) Heatmap of the top 20 most highly induced genes at 5, 10, and 30 min after touch. Genes with peak induction at 5 min that maintain high expression through 30 min are highlighted in green, (e) PCA of pooled transcriptome profiles across a 0-180 min time course.

Among the total genes identified from differential expression analysis, 1,574 genes were found to be responsive to touch (Fig. S1; Table S4). Overall, touch predominantly altered expression through up-regulation at all time points (Fig. S1). DEGs induced at 5 minutes generally exhibited sustained up-regulation over the subsequent 30-minute window (Table S4). In particular, transcription factors (TFs) play a key role in mediating early touch signaling and transducing signals to a wide range of downstream touch-responsive gene sets. The dramatic changes observed at 5 minutes post-touch highlight the importance of transcriptional regulation in touch signaling (Fig. S1; Table S5). Targeted analysis revealed a significant enrichment of TFs among DEGs at each time point following touch, with 47 (21.6% of DEGs at 5 min), 96 (14.2% of DEGs at 10 min), and 166 (11.7% of DEGs at 30 min) TFs identified, respectively (Fig. 1b). The proportion of TF-encoding genes among DEGs decreased over the 30-minute period. Although the absolute number of identified TFs varied across time points, ETHYLENE RESPONSE FACTOR (ERF) families exhibited significant enrichment at all time points post-touch. The fold enrichment of ERF TFs decreased over time, suggesting their prominent role in early mechanical signaling (Fig. 1c). In contrast to ERF TFs, WRKY TFs showed significant enrichment only at 10 minutes post-touch. This pattern differs from that observed in sorghum subjected to leaf removal, where WRKY TFs were enriched at both 10 and 30 minutes following treatment (Fig. 1c; Park et al., 2024). This suggests that ERF and WRKY TFs may play a role in early mechanical responses within 30 minutes of mechanical stimulation.

To further verify TFs potentially crucial for the rapid touch response, the TF2Network tool (Kulkarni et al., 2018) was used to predict transcription factor binding sites within the up-regulated DEGs at each time point. Position weight matrix (PWMs) mapping showed significantly enriched binding sites for CAMTA, WRKY, and MYC TF families in the up-regulated DEGs across all time points tested by this experiment, highlighting their importance in mechanical responses (Table S6; Van Moerkercke et al., 2019; Darwish et al., 2022). Binding sites for a wide variety of ERF TF families were identified in the touch-responsive up-regulated DEGs at 10 and 30 minutes after touch (Table S6a-c). The top 20 most induced *Arabidopsis* genes at each time point post-touch were examined to identify those most highly responsive to mechanical stimulation. In particular, *DDF1*, *AT4G27654*, *CBF4*, and *RRTF1* showed a strong initial response at 5 minutes and sustained high expression through 30 minutes (Fig. 1d). Given that *DDF1*, *CBF4*, and *RRTF1* belong to the ERF family, this finding aligns with the observation that ERF TF members were significantly enriched across all time points post-touch in this experiment.

The long-term temporal transcriptional response to touch remains poorly characterized. To address this, we compiled a comprehensive dataset. This dataset integrated RNA-seq data generated in this study with publicly available RNA-seq profiles of *Arabidopsis* in response to touch or water spray (Table S7; Xu et al., 2019; Wang et al., 2023). Principal component analysis (PCA) of read counts transformed by variance stabilizing transformation (VST), with batch correction applied, showed that biological replicates from different experiments clustered closely within their respective time points (Fig. 1e). Interestingly, samples from 180 minutes post-touch clustered closely with basal (control) samples, suggesting that most touch-induced transcriptional changes are resolved or terminated by 180 minutes (Fig. 1e). While our study and the transcriptome data from Wang et al. (2023) used 11- to 15-day-old plants, Xu et al. (2019) analyzed four-week-old *Arabidopsis* plants. This implies that the overall transcriptional changes in response to touch are consistent across this range, despite the variation in plant age.

To identify potential key regulators of early mechanical stimulation, we focused on genes identified in the pooled data that were rapidly up-regulated at 5 minutes post-touch, maintained high expression levels for 40-60 minutes, and subsequently decreased. *RRTF1*, *AT4G27654*, and *DDF1* were potential candidates. Among these candidates, RRTF1 was prioritized as a key candidate due to its characteristic rapid spike and decay in expression. A conserved role for RRTF1 involving early mechanical responses has been demonstrated across several species, ranging from *Arabidopsis* to cereal crops like sorghum and wheat (Li et al., 2023; Darwish et al., 2023; Park et al., 2024). The temporal expression pattern of *RRTF1* typically showed a rapid decrease within 60 minutes post-stimulation, a trend observed irrespective of the species. Notably, this expression pattern of *RRTF1* is observed not only in response to gentle touch but also following more severe stimuli like leaf removal (a form of mechanical stimulation). Collectively, these observations led to the hypothesis that *RRTF1* may be a key regulator of early mechanical responses.

### No significant role of *RRTF1* in major touch-induced thigmomorphogenic responses to gentle touching

Prolonged and repeated touch has been shown to elicit a wide range of morphological changes in plants (Braam and Davis, 1990; Braam, 2005; Chehab et al., 2012). Given the rapid and significant induction of *RRTF1* in response to touch, we hypothesized its potential role in thigmomorphogenesis. To genetically investigate whether *RRTF1* plays a role in thigmomorphogenesis, two t-DNA mutants in the *AT4G34410* locus (*rrtf1-1*; SALK_ 206786C and *rrtf1-2*; SALK_150614) were obtained and their touch-induced morphological changes were assessed (Fig. S2). Both lines have insertions approximately two-thirds into the coding sequence that disrupt the DNA-binding domain, with the insertions oriented in opposite directions. Touch-induced morphogenic responses can vary depending on mechanical load, frequency of stimulation, and overall plant growth conditions. The most representative thigmomorphogenic responses include reduced plant shoot height and delayed flowering (Wang et al., 2018; Darwish et al., 2022; Wang et al., 2023). Repetitive touch can also alter leaf morphology, leading to reduced rosette area and petiole length (Garner and Langton, 1997; Liu et al., 2007; Chehab et al., 2012; Wang et al., 2023).

10-day-old, soil-grown seedlings were subjected to gentle touching (10 times) with a soft paintbrush twice daily until the onset of flowering for each plant. Brush-touched wild-type (Col-0) plants exhibited a clear delay in flowering time, approximately 1.5 day later compared to untouched Col-0 control plants. A similar delay in flowering (approximately 1.2-1.7 days relative to their respective untouched controls) was observed in brush-touched *rrtf1* mutant (Fig. S2b). Contrary to expectations, both wild-type and *rrtf1* mutant exhibited delayed flowering, with no significant difference between the genotypes under these conditions. Other phenotypes of brush-touched plants at flowering were nearly identical to that of untouched control plants regardless of genotype (Fig. S2c-f). For example, brush touching had no significant impact on rosette area at the time of flowering in either wild-type or *rrtf1* mutant plants under these conditions (Fig. S2c).

To determine whether repetitive touch had any further impact on plant development after flowering, wild-type and *rrtf1* mutant plants were grown for an extended period and harvested 6 days after flowering. The impact of mechanical stimulation on plant architecture has been documented across various species; for instance, touch treatment reduces branch diameter in potato (Markovic et al., 2016), while wind exposure modulates the degree of branching in *Arabidopsis* (Pigliucci, 2002). Prolonged brushing reduced the length of the first three rosette branches in Col-0 and the *rrtf1* mutant compared to untouched control plants, whereas the frequency of rosette bud outgrowth remained unaffected by touch treatment (Fig. S2e,f).

### Attenuated thigmomorphogenesis in *rrtf1* mutant under aggressive mechanical stimulation

Our initial gentle touch treatment, using an intensity comparable to some previously described systems, did not consistently or fully induce all characteristic thigmomorphogenic responses in wild-type (Col-0) plants. Although reduced rosette leaf area is a representative thigmomorphogenic response (Jaffe, 1973; Chehab et al., 2012; Graham and Wheeler, 2017; Darwish et al., 2022; Wang et al., 2023), our gentle touch conditions had no significant effect on rosette leaf area at flowering in any of the genotypes (Fig. S2c). Consistent with this, while mechanically stimulated plants can exhibit reduced shoot fresh weight under certain growth conditions (Jaffe, 1973; Biddington, 1985; Kläring, 1999; Graham and Wheeler, 2017; Gladala-Kostarz et al., 2020; Hindhaugh et al., 2021), the gentle touch treatment in our study had no impact on shoot fresh weight and branching (Fig. S2d).

These observations highlight that the manifestation of thigmomorphogenesis is highly dependent on plant growth conditions and the specifics of the mechanical treatment. To elicit more robust and consistent touch-induced phenotypic changes, we developed an automated, aggressive touch treatment that allowed for increased frequency and duration of mechanical stimulation (Video S1). Given the expression characteristics of the *RRTF1* gene observed in our pooled RNA-seq analysis, we devised an optimized protocol for aggressive mechanical stimulation (Fig. S3). *RRTF1* is typically activated within 5 minutes of touch, with expression levels subsequently diminishing, usually within 40 to 60 minutes post-stimulation. To promote sustained *RRTF1* expression, the aggressive touch protocol applied three rounds of stimulation per day at 8-hour intervals. Each round consisted of 40 touches over the course of 1 hour. The aggressive touch experiment induced conspicuous thigmomorphogenic changes, including reduced rosette leaf area and shoot height, and delayed flowering (Fig. 2a-d). In contrast to the gentle touch treatment using a paintbrush, aggressive mechanical stimulation significantly altered leaf morphology, including rosette leaf area and petiole length (Fig. 2c; Fig. S3b). To quantitatively compare WT and *rrtf1* responses, we calculated the percentage of touch-induced relative changes for each parameter. Across all thigmomorphogenic parameters collected, focusing on shoot height as an example, touch reduced wild-type plant height to 37-64 % of their untouched controls, whereas *rrtf1-1* mutant exhibited an attenuated response, growing to 46-93 % of their corresponding untouched controls (Fig. 2e). The subtle but consistent attenuated height reduction observed in *rrtf1* mutant under aggressive touch treatment suggest that RRTF1 promotes touch-induced height reduction in *Arabidopsis*.

**Fig. 2.**
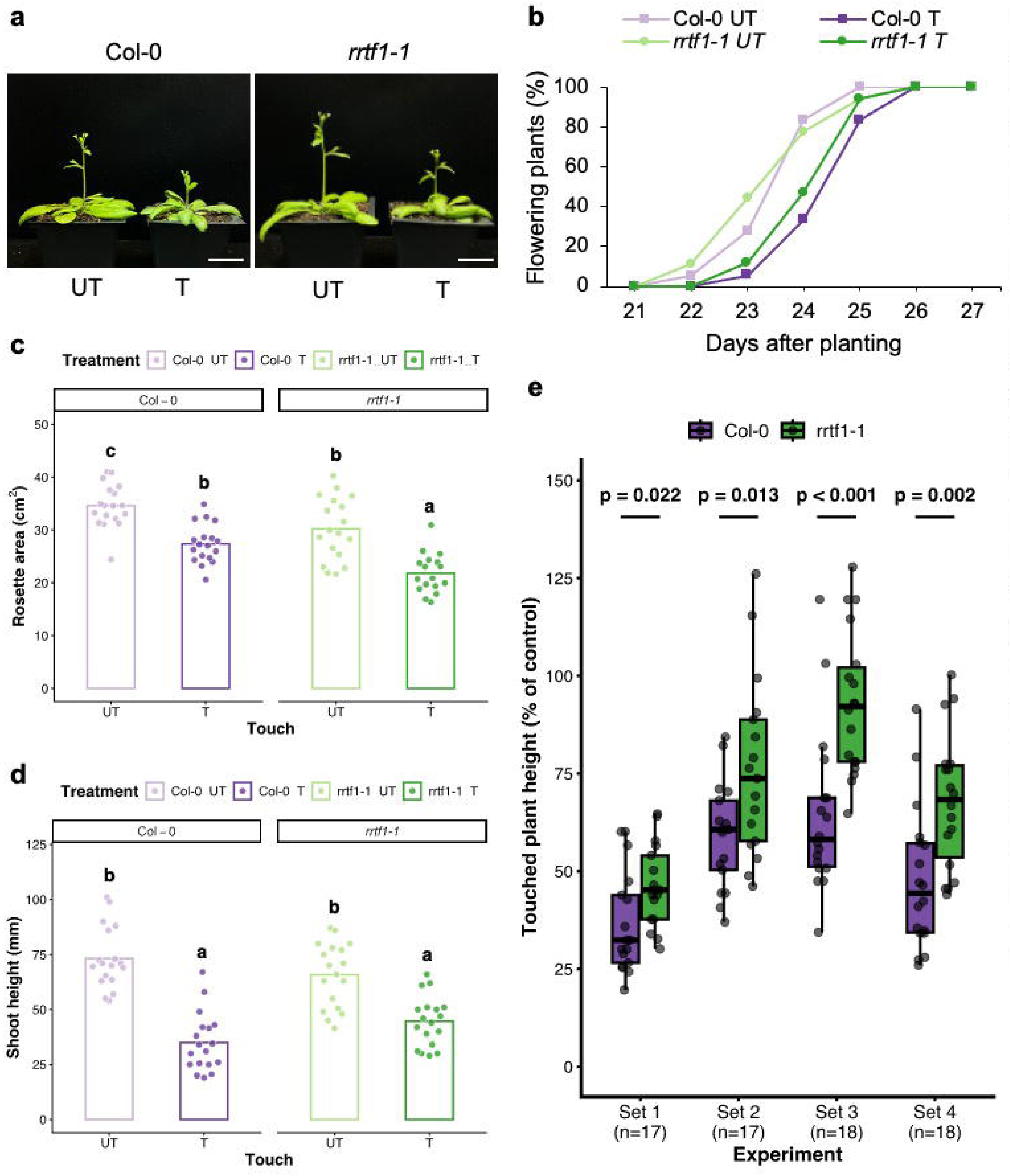
Attenuated thigmomophogenesis in the *rrtf1* mutant under aggressive mechanical stimulation, (a) Photographs of Col-0 and *rrtf1-1* mutant at the time each individual plant began to flower. UT and T indicate the untouched (control) and touched groups, respectively. Bar, 2 cm. (b) Percentage (%) of flowering plants following aggressive touch over the growth period (days after planting), (c, d) Rosette leaf area (c) and shoot height (d) of Col-0 and *rrtf1* mutant in response to aggressive touch, measured at the onset of flowering for each plant, (e) Shoot height of touched plants relative to their corresponding untouched controls across four independent experiments (Sets 1-4). Statistical significance was determined by two-way ANOVA followed by Tukey’s post-hoc tests. Different letters indicate significant differences among all groups at *p* < 0.05.

### Attenuated thigmomorphogenesis in *rrtf1* mutant cannot be rescued by exogenous JA application

Jasmonic acid (JA), a key regulator of plant growth and defense, plays a critical role in mechanical signaling pathways. Previous studies have shown that not only *Arabidopsis* mutants defective in JA biosynthesis and signaling exhibit impaired thigmomorphogenesis (Chehab et al., 2012; Van Moerkercke et al., 2019), but JA is rapidly induced within 2.5 minutes after leaf removal in sorghum (Park et al., 2024). Moreover, previous studies have characterized the crosstalk between RRTF1 and JA signaling pathways. The basic helix-loop-helix transcription factor MYC2, a key regulator of JA signaling, directly activates *RRTF1* expression (Van Moerkercke et al., 2019); accordingly, the *myc2 myc3 myc4 myc5* quadruple mutant showed a remarkably reduced *RRTF1* induction 22 minutes after touch compared to the wild-type (Darwish et al., 2022). RRTF1 is also implicated in JA-dependent wound regeneration (Zhang et al., 2019). In wild-type seedlings, *RRTF1* transcripts peaked (∼20-fold) 20 minutes after methyl jasmonate (MeJA) application and returned to baseline within 1 hour (Cai et al., 2014; Hickman et al., 2017).

We confirmed that MeJA elicited efficient *RRTF1* induction within 15 minutes, even in the absence of mechanical stimulation (Fig. 3a). Prior studies showed that JA levels in sorghum buds and stems are highly responsive to mechanical stress (Park et al., 2024). To validate that mechanical stimulation induces JA accumulation in *Arabidopsis*, we quantified endogenous phytohormone levels in wild-type and *rrtf1* mutant plants before and after touch with a paintbrush (Fig. 3b; Fig. S4). Endogenous JA levels in *Arabidopsis* rapidly increased within 10 minutes of touch treatment in both wild-type and *rrtf1* mutant plants, consistent with previous findings in sorghum subjected to leaf removal, a more aggressive form of mechanical stimulation (Fig. 3b; Park et al., 2024). The overall JA responsiveness to touch was similar between wild-type (Col-0) and *rrtf1* mutant. However, *rrtf1* mutant showed a slightly faster JA response to touch, suggesting a potential modulatory role for *RRTF1* in early JA signaling pathways. For other phytohormones, indole-3-acetic acid (IAA), abscisic acid (ABA), and bioactive gibberellin (GA_4_) levels remained largely unchanged during the first 10 minutes of touch in both wild-type and the *rrtf1* mutant (Fig. S4). However, basal levels of GA precursors such as GA_5_ and GA_20_ were higher in *rrtf1* mutant compared to the wild-type, whereas basal levels of the bioactive GA_1_, GA_4_, and the inactive GA_8_ were similar between the genotypes. These findings suggest that while *rrtf1* loss-of-function mutation elevates basal levels of GA precursors, this increase does not translate into different levels of bioactive GA_1_ and GA_4_ between genotypes during the 10-minute touch treatment.

**Fig. 3.**
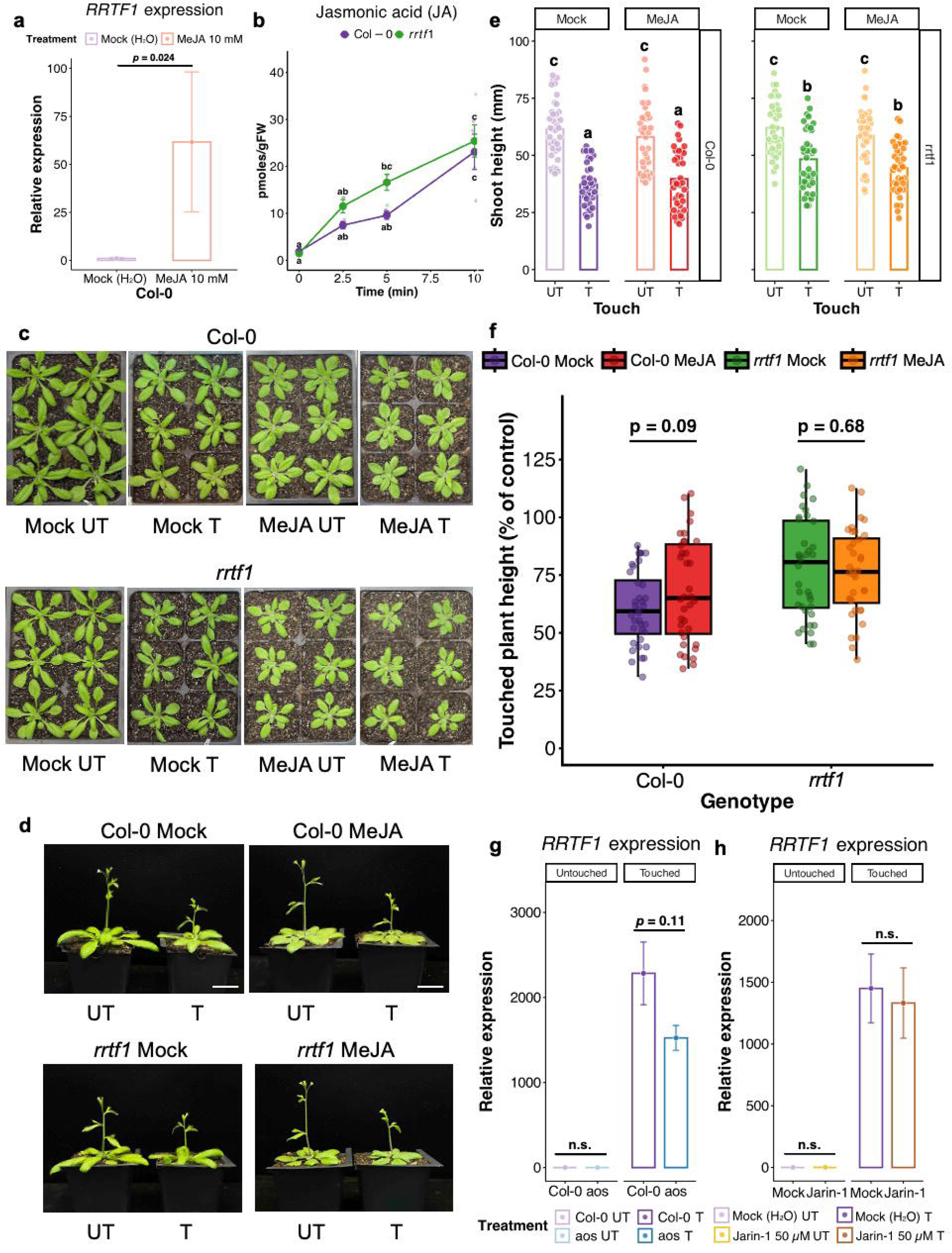
Attenuated thigmomorphogenesis in the *rrtf1* mutant cannot be rescued by exogenous JA application, (a) Relative transcript levels of *RRTF1* in Col-0 under 10 mM MeJA treatment. The y-axis represents fold change relative to the untouched (0 min) control (set to 1). (b) Jasmonic acid (JA) abundance in Col-0 and *rrtf1* mutant at 0, 2.5, 5, and 10 min after touch, (c) Photographs of Col-0 and *rrtf1* mutant at the time the first plant began to flower (22 days after planting) under mock (water) and 50 μM methyl jasmonate (MeJA) treatments, (d) Photographs of Col-0 and *rrtf1* mutant at the time each individual plant began to flower (considered ready for harvest), (e) Shoot height of Col-0 and *rrtf1* mutant following aggressive touch and 50 μM MeJA treatment, measured at the onset of flowering for each plant, (f) Shoot height of touched plants relative to their corresponding untouched controls under mock and 50 μM MeJA treatments, (g) Relative transcript levels of *RRTF1* in Col-0 and *aos* mutant after touch. The y-axis represents fold change relative to untouched (0 min) Col-0 control (set to 1). (h) Relative transcript levels of *RRTF1* in Col-0 at 10 min after touch, treated with either mock (water) or 50 μM Jarin-1. The y-axis represents fold change relative to the mock-treated, untouched Col-0 control (set to 1). Statistical significance was determined by two-way ANOVA followed by Tukey’s post-hoc tests. Different letters indicate significant differences among all groups (p < 0.05). UT and T indicate the untouched (control) and touched groups, respectively. Bar, 2 cm

To investigate whether JA impacts *RRTF1*-mediated thigmomorphogenesis over time, 50 μM MeJA was applied to wild-type and *rrtf1* mutant. Consistent with the established role of JA in thigmomorphogenesis, exogenous MeJA treatment induced substantial alterations in leaf morphology (Fig. 3c). However, these responses remained phenotypically distinct from those elicited by mechanical stimulation in both wild-type and *rrtf1* mutant. Specifically, MeJA-treated plants exhibited rounder and flatter leaves, characterized by a lower ratio of leaf blade length to width, regardless of genotype or touch treatment (Fig. S5a). The combination of MeJA and touch treatment resulted in the most significant reduction in rosette leaf area in both wild-type and *rrtf1* mutant (Fig. S5b). These findings suggest that JA has additive roles in certain aspects of thigmomorphogenesis, acting via a *RRTF1*-independent pathway for these particular leaf phenotypes.

However, flowering time exhibited a genotype-specific response to MeJA. In the wild-type, MeJA treatment abolished the touch-induced delay in flowering, whereas the *rrtf1* mutant flowered later than the wild-type under MeJA treatment (Fig. S5c). MeJA had minimal impact on touch-induced shoot height reduction in either wild-type or *rrtf1* mutant compared to mock treatment (Fig. 3e). Similarly, the percentage height of touched plants relative to untouched plants was unaffected by MeJA, regardless of RRTF1 genotype (Fig. 3f). These results suggest that touch-induced height reduction is regulated by RRTF1, and the *rrtf1* mutant phenotype of attenuated height reduction cannot be rescued by exogenous JA.

Despite the well-known connections between RRTF1 and JA signaling, we found that JA accumulation following touch treatment in *rrtf1* mutant was essentially indistinguishable from the wild-type (Fig. 3b). To investigate whether JA is required for *RRTF1* induction by touch, we assessed the expression of *RRTF1* in response to mechanical stimulation in JA-deficient *aos* (*allene oxide synthase*) mutant. Touch treatments triggered *RRTF1* transcription in *aos* mutant to the nearly same extent as in wild-type (Fig. 3g). Similarly, the suppression of JA biosynthesis by Jarin-1, a JA-Ile biosynthesis inhibitor, did not compromise *RRTF1* expression in response to touch (Fig. 3h). These results suggest that the initial induction of *RRTF1* occurs largely independently of JA biosynthesis in early touch signaling.

Given that both touch-induced expression of the JA signaling component *JAZ7* (*JASMONATE-ZIM-DOMAIN PROTEIN 7*) and JA accumulation in the *rrtf1* mutant were comparable to the wild-type (Fig. 3b; Fig 4b), and that *RRTF1* induction by touch occurs independently of JA biosynthesis (Fig. 3g,h), it is likely that RRTF1 regulates the height response to touch via a JA-independent pathway. Together, these results suggest that RRTF1 and JA pathways converge additively on leaf thigmomorphogenic responses, whereas RRTF1 acts independently of JA in regulating touch-mediated shoot height reduction.

**Fig. 4.**
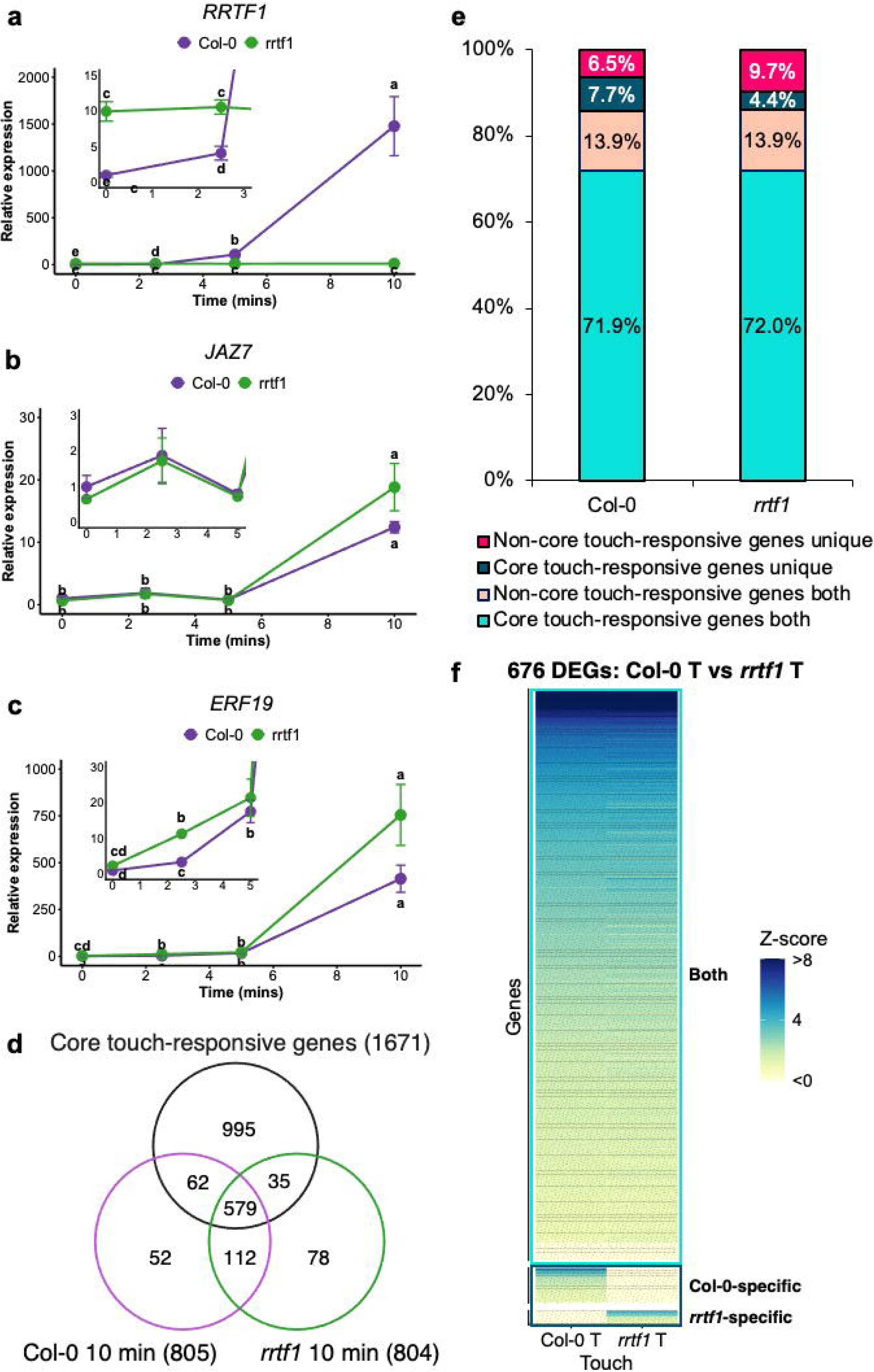
The core touch-responsive transcriptome is retained in the *rrtf1* mutant, (a-c) Relative transcript levels of *RRTF1* (a), *JAZ7(b),* and *ERF19* (c) in Col-0 and *rrtf1* mutant after touch. Insets show the first 2.5-5 min of the time course. The y-axis represents fold change relative to the untouched (0 min) Col-0 control (set to 1). (d) Venn diagram showing the overlap of differentially expressed genes (DEGs) in Col-0 and *rrtf1* mutant at 10 min post-touch with previously identified core touch-responsive genes, (e) Distribution of DEGs in Col-0 and *rrtf1* mutant at 10 min post-touch. Genes are categorized as core or non-core touch-responsive, and further subdivided into those commonly expressed in both genotypes versus those uniquely expressed in either Col-0 or *rrtf1* mutant, (f) Heatmap of core touch-responsive genes from our transcriptome data. Each row represents an individual gene, and the color scale indicates scaled expression (z-score). Boxes highlight commonly expressed genes and genotype-specific genes, respectively.

### The core touch-responsive transcriptome is retained in the *rrtf1* mutant

To examine how quickly early gene expression, such as *RRTF1,* responds to touch, we performed RT-qPCR on wild-type and *rrtf1* mutant at 2.5, 5, and 10 minutes post-stimulation. Based on our transcriptome analysis and prior literature, we selected RRTF1 and other signaling markers known for their robust transcriptional responses. The *RRTF1* expression in wild-type plants increased within 2.5 minutes after touch (Fig. 4a), consistent with its pattern in the *Arabidopsis* transcriptome datasets. The *RRTF1* expression in *Arabidopsis* exhibited a pattern similar to that observed in sorghum following leaf removal (Park et al., 2024). To determine if *RRTF1* influences the expression of other touch-responsive genes, *JAZ7* and *ERF19* were selected for analysis in the *rrtf1* mutant background. *JAZ7* was previously reported to be induced by mechanical stimulation within 15 minutes (Wang et al., 2018). *JAZ7* transcript levels were induced in both wild-type and *rrtf1* mutant plants by 10 minutes post-touch (Fig. 4b). We also found that *ERF19*, which was previously shown to be expressed 25 minutes after water spray (Van Moerkercke et al., 2019), was induced within 2.5 minutes of touch in our system. Although *ERF19* appeared to show a slightly faster or stronger initial response in the *rrtf1* mutant at very early time points, its overall expression trend within the first 10 minutes was comparable between both genotypes (Fig. 4c). Similarly, the stress-responsive gene *CBF2*, a member of the ERF TF family, exhibited comparable transcript levels in both wild-type and *rrtf1* mutant plants at 10 minutes post-touch (Fig. S6). Collectively, although *RRTF1* itself showed significant transcriptional induction after touch in wild-type plants, these findings suggest that *RRTF1* may not be directly involved in regulating the expression of these previously known touch-responsive genes (*JAZ7*, *ERF19*, and *CBF2*) at the transcriptional level during the early response to a single touch event.

We then profiled touch-induced global transcriptomes of wild-type and *rrtf1* mutant to define how RRTF1 regulates downstream touch-responsive gene expression. We selected the 10-minute time point as *RRTF1* expression typically peaks at this time. Plants were gently brushed 15 times and harvested 10 minutes after the single treatment. Untouched plants were harvested concurrently to control for any stimulation caused by the harvesting process. To identify differentially expressed genes (DEGs) associated with touch in wild-type and *rrtf1* mutant backgrounds, cleaned reads (approximately 99% of the 20-57 million acquired per sample) were mapped to the *Arabidopsis* genome (TAIR10, Lamesch et al., 2012). A total of 20,840 genes were identified as expressed, based on the criterion of having raw counts > 10 in at least three samples. A PCA of the variance stabilizing transformation (VST)-normalized read counts separated the effect of the touch treatment (Fig. S7a). Touch treatment altered gene expression predominantly through up-regulation in both genotypes. The number of DEGs was similar between wild-type and *rrtf1* mutant at 10 minutes after touch: 805 and 804 DEGs, respectively (Table S8).

To further elucidate the role of RRTF1 in the genome-wide transcriptional response to touch, we compared the transcriptomes of touched wild-type and *rrtf1* mutant against untouched wild-type (Fig. 4d). In total, 804 genes showed an altered touch response in the *rrtf1* mutant, while 805 genes had an altered touch response in wild-type (FDR < 0.05, >1.3-fold change, determined using edgeR glmTreat). While the number of touch-responsive genes was similar, the specific transcriptome profiles had distinct elements (Fig. 4d). 691 genes were commonly induced 10 minutes following touch in both genotypes (Table S12). A relatively large portion of these common genes (579 out of 691) belonged to the previously defined set of 1,671 “core touch-responsive” genes, which are commonly differentially expressed across a wide range of mechanical stimulation transcriptome (Van Moerkerke et al., 2019). A majority of touch-induced genes, whether or not core touch-responsive, were commonly induced in both wild-type and *rrtf1* mutant (Fig. 4e). Overall, 676 core touch-responsive genes showed differential expression in response to touch, and we plotted the expression values of these genes in heatmaps (Fig. 4f). The commonly expressed core touch-responsive genes showed similar expression trends between wild-type and the *rrtf1* mutant in response to touch. Of the 242 non-core touch-responsive genes considered, wild-type and the *rrtf1* mutant commonly expressed 112 genes in response to touch (Fig. 4d). This comparison suggests that the touch responses of wild-type and *rrtf1* mutant at 10 minutes after touch are largely similar. Approximately 86% of the touch-responsive DEGs identified in wild-type were also present in the *rrtf1* mutant, showing similar expression patterns. Consequently, the remaining 14% of DEGs constitute candidate genes that likely underlie the differential thigmomorphogenic responses observed between the two genotypes.

### RRTF1 modulates a subset of touch-responsive WRKY-dependent regulon

Gene Ontology (GO) Molecular Function (MF) enrichment differed markedly between the wild-type-specific and *rrtf1*-specific touch-induced gene sets (Table S10). To identify candidate upstream regulators, these unique gene sets were further analyzed for overrepresented transcription factor binding sites using TF2Network (Kulkarni et al., 2018). In total, 698 motifs were identified as overrepresented in the promoters of commonly up-regulated genes in both wild-type and *rrtf1* mutant (Table S11a). A variety of DNA motifs were overrepresented among these common genes, including binding sites for CAMTA, WRKY, and ERF TFs (Table S11a). The CAMTA, WRKY, and ERF TFs have been previously proposed to be involved in mechanical stress responses by activating downstream signaling pathways (Darwish et al., 2022; Matsumura et al., 2022; Li et al., 2023; Park et al., 2024).

When examining the DEGs unique to each genotype, WRKY and NAC binding motifs were overrepresented in the wild-type-specific DEGs, whereas bZIP and bHLH motifs were enriched in the *rrtf1*-specific gene set. Interestingly, WRKY motifs were no longer overrepresented among the *rrtf1*-specific DEGs. This implies that WRKY-dependent activation is compromised in the absence of RRTF1 (Fig. 5a, Table S11b,c). To further validate these findings, we conducted promoter motif analysis comparing touch-induced genes specific to Col-0 with those specific to the *rrtf1* mutant. Promoter regions of wild-type specific genes contained an AGTCAAC motif, which is a known WRKY-binding motif (Fig. 5b; Pan et al., 2009). WRKY40, along with its closely related paralogs WRKY18 and WRKY60, has been implicated in mechanical responses (Darwish et al., 2022; Matsumura et al., 2022). The *rrtf1* mutant showed wild-type like expression of WRKY TFs after touch (Fig. S7b). Similar uncoupling is observed in other mechanosensory mutants: the *wrky18 wrky40 wrky60* triple mutant showed wild-type like *RRTF1* expression at 22 minutes after touch (Darwish et al., 2022). Collectively, these findings suggest that RRTF1 might collaborate with WRKY TFs to drive the expression of a distinct subset of touch-responsive genes, without altering WRKY transcript abundance itself.

**Fig. 5.**
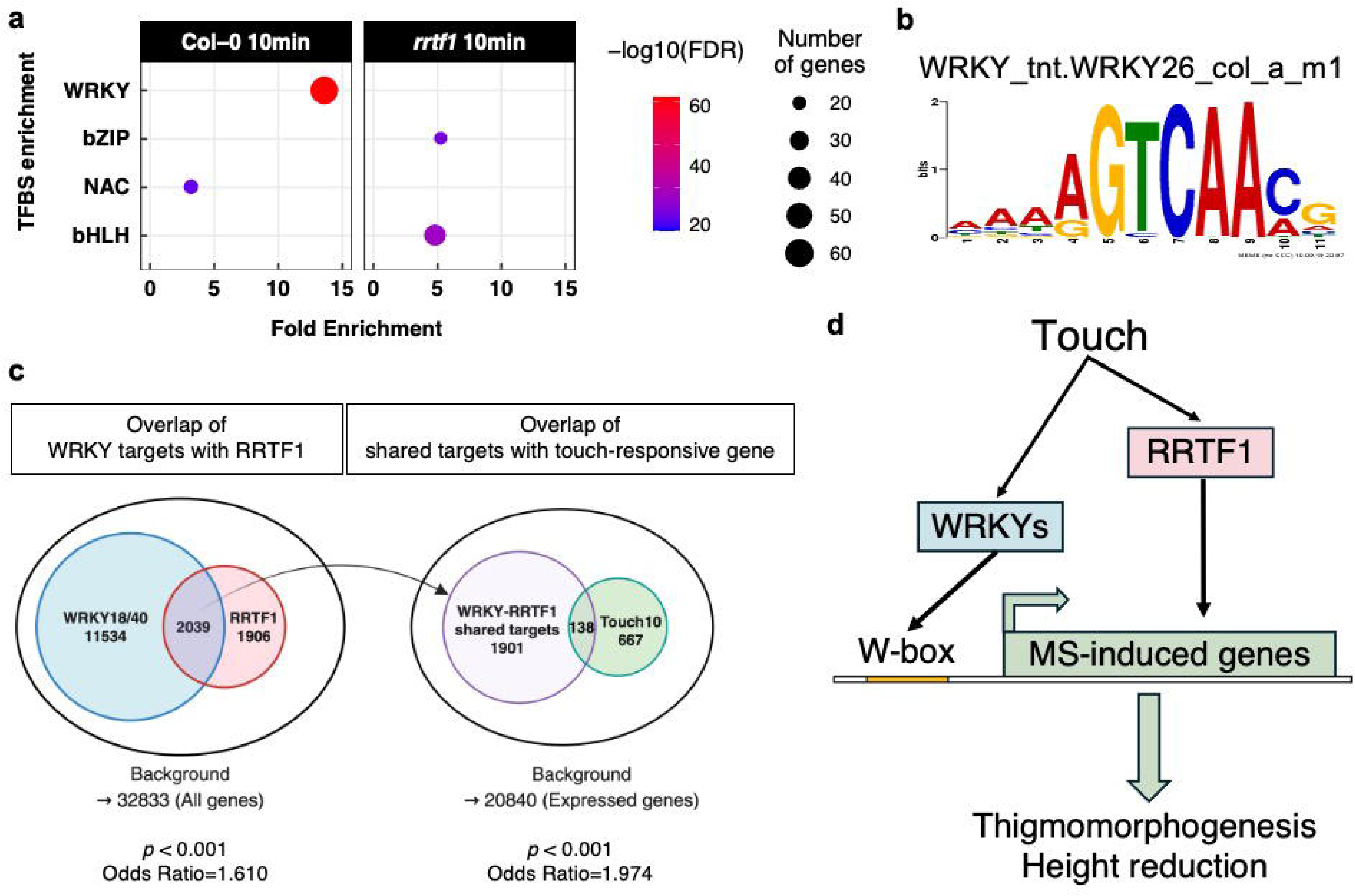
Modulation of touch-responsive WRKY-dependent regulons by RRTF1. (a) Transcription factor binding site enrichment analysis of commonly and uniquely DEGs in Col-0 and *rrtf1* mutant, (b) Transcription factor binding motifs overrepresented in touch-responsive genes specific to Col-0 compared with those specific to the *rrtf1* mutant, (c) Overlap of DEGs between RRTF1 and WRKY18/40 regulons within the touch-responsive transcriptome using DAP-seq binding profiles of RRTF1 and WRKY18/40. (d) A working model illustrating the roles of RRTF1 and WRKY transcription factors in thigmomorphogenesis.

To investigate the interplay between RRTF1 and WRKY transcription factors, we determined the binding profiles of RRTF1 and WRKY18/40 using DAP-seq (O’Malley et al., 2016). Approximately 52% of RRTF1 target genes overlapped with WRKY18/40 targets (*p* < 0.001), indicating a potential regulatory link (Fig. 5c). In addition, we analyzed the intersection of these shared targets with touch-responsive genes from our experiment. Notably, touch-responsive genes were significantly enriched within the RRTF1-WRKY18/40 shared target set (*p* < 0.001), suggesting a regulatory network connecting RRTF1 and WRKY18/40 to the transcriptional touch response (Fig. 5c).

## Discussion

### Temporal dynamics of transcriptional reprogramming during the touch response

Mechanical stimulation initiates a rapid, multilayered signaling cascade that ultimately remodels plant growth and development. In this study, our large-scale RNA-seq analysis, spanning 5 to 180 minutes post-stimulation, revealed that 9,771 genes-approximately 30% of the *Arabidopsis* genome-are differentially expressed in response to mechanical stimulation (Table S7). While touch-induced phenotypic alterations are well documented across diverse plant species, the precise chronological orchestration of the earliest signaling events has remained elusive. Our data demonstrate that this transcriptional rewiring is exceptionally rapid. Within just 5 minutes of stimulation, an initial wave of ∼200-300 genes, enriched for transcriptional regulation, defense, and ethylene signaling, is rapidly induced (Fig. 1b; Fig. S1b; Wang et al., 2023). Few studies have resolved how dynamically these transcriptional profiles shift across such brief intervals.

### Connecting *RRTF1* transcription to touch-induced developmental plasticity

While several studies across different plant species have demonstrated that *RRTF1* is rapidly induced by various mechanical stimuli, such as water spray and leaf excision, its specific developmental function in thigmomorphogenesis has remained largely unexplored. Although *RRTF1* is known to mediate plant defense responses (Li et al., 2021; Singh et al., 2023), its contribution to mechanosensitive phenotypic plasticity is undefined. To elucidate the precise role of *RRTF1* in thigmomorphogenesis, we employed a controlled mechanical stimulation system capable of delivering various intensities of mechanical stress (Fig. S3). By evaluating plant responses across distinct intensities of mechanical stimulation, we quantified specific, *RRTF1*-dependent thigmomorphogenic phenotypes, thereby linking early transcriptional dynamics to the ultimate morphological adaptation. Thigmomorphogenesis depends on the intensity of the mechanical stimulation (Hunt and Jaffe, 1980; Kläring, 1999). When subjected to mild mechanical stimulation (gentle brushing 10-20 times, twice daily), the *rrtf1* mutant exhibited thigmomorphogenic responses comparable to the wild-type, including delayed flowering, altered branching, and reduced stem height and rosette leaf area. Only when mechanical load was intensified (high-frequency, triple daily), *rrtf1* mutant plants exhibited an attenuated reduction in stem height. Thus, *RRTF1* activates growth inhibition under chronic or stronger mechanical stress, acting as a gain controller rather than an on/off regulator of thigmomorphogenesis.

Given that early touch-responsive gene expression typically diminishes within 60 minutes post-touch stimulation, our devised methods for quantifiable and repeatable thigmomorphogenesis maintained an active physiological state for approximately 2 hours of touch responses (i.e., a 60-minute response period following each of the three daily sessions). Our data suggest that under severe or persistent mechanical stress, a key function of *RRTF1* is to promote the thigmomorphogenic response (Fig. 2, Fig. S2). This positions *RRTF1* as a component that helps fine-tune the plant’s morphological adaptation to the perceived level of mechanical stimulation. In agricultural fields, rain and gust conditions can last for hours, and densely planted crops are constantly in physical contact with their neighbors. *RRTF1* may be an important player for modulating crop growth in the field, especially in dense planting conditions.

### Compensatory networks maintain early touch signaling in *rrtf1* mutant

Transcriptome profiling of the *rrtf1* mutant at 10 minutes post-touch revealed a global transcriptional landscape highly comparable to that of wild-type plants (Fig. 4e, Fig. S7a). Approximately 86% of touch-responsive differentially expressed genes (DEGs) were shared between both genotypes, displaying similar transcriptional dynamics. These findings indicate that *RRTF1* does not act as a singular master regulator for the early touch transcriptional response. Instead, they highlight an alternative transcriptional network capable of mounting a robust response even in its absence.

We performed GO enrichment analysis on the ∼14% of DEGs that were uniquely or differentially regulated between the wild-type and *rrtf1* mutant post-touch (Fig. S7c, Table S9). Interestingly, an evaluation of the top 10 significantly enriched GO terms revealed considerable overlap between the two genotype-specific upregulated gene sets. For example, the broader “oxylipin biosynthetic process” was significantly enriched in the unique DEGs of both genotypes. However, this functional convergence was driven by the specific induction of *OPR1* (*12-OXOPHYTODIENOATE REDUCTASE 1)* in the wild-type, contrary to the unique upregulation of *LOX5* (*LIPOXYGENASE 5*) in the *rrtf1* mutant (Table S9).

### Complex crosstalk between RRTF1 and JA in the touch response

Our investigation into the interplay between RRTF1 and JA signaling uncovers a highly complex regulatory network. While molecular crosstalk between these pathways is well documented, such as the physical interaction between RRTF1 and JAZ9 (Zhang et al., 2019) and the insensitivity of *rrtf1* mutant to JA-promoted root hair elongation (Sui et al., 2025), our data demonstrate that the initial touch-induced activation of *RRTF1* occurs independently of JA. Although *RRTF1* expression is inducible by exogenous MeJA via MYC transcription factors (Van Moerkercke et al., 2019), its rapid upregulation following mechanical stimulation remained largely unperturbed in the *aos* mutant and under Jarin-1 inhibitor treatment (Fig. 3g,h). Nonetheless, the attenuated height reduction observed in the *rrtf1* mutant under aggressive mechanical stimulation persisted regardless of MeJA treatment (Fig. 3f).

### RRTF1-WRKY regulatory module fine-tunes thigmomorphogenesis

The loss of WRKY TF binding site enrichment among the *rrtf1*-specific DEGs, despite *WRKY* gene expression itself being largely unaffected in the *rrtf1* mutant, points to a cooperative regulatory relationship (Fig. 5a, Fig. S7b). Since the activation of specific WRKY-dependent targets requires RRTF1 (Fig. 5c), RRTF1 likely acts as a critical co-regulator, fine-tuning the expression of regulons essential for the broader mechanosensory response.

In conclusion, here, we demonstrate that RRTF1 mediates specific morphological adaptations via a pathway that functions independently of canonical JA signaling. Rather than acting as a universal on/off switch for early touch responses, RRTF1 emerges a modulatory node that specifically co-regulates subsets of touch-responsive genes through the RRTF1-WRKY module (Fig. 5d). Elucidating these signaling networks provides a molecular framework with significant implications for modern agriculture, particularly where high-density planting results in frequent physical contact between neighboring plants and consequent thigomorphogenesis. By uncovering the genetic mechanisms underlying thigmomorphogenesis, this study highlights potential targets for engineering or breeding crops that retain robust growth, despite frequent mechanical stimulation, without yield penalties.

## Materials and Methods

### Plant material and growth conditions

The ecotype Columbia (Col-0) of *Arabidopsis* (*Arabidopsis thaliana*) was used throughout. The *rrtf1-1* (SALK_206786C), *rrtf1-2* (SALK_150614), and *aos* (SALK_017756) seeds were obtained from the *Arabidopsis* Biological Resource Center (Ohio State University, Columbus). The mutant used in the study was genotyped using standard PCR protocols to confirm T-DNA insertions and homozygosity.

Seeds were stratified at 4°C for 3 days and sown in six-cell inserts filled with a custom media mixture (36 plants per tray) supplemented with 2mL of 1x Hoagland’s solution. A custom media mix was prepared by combining peat moss, vermiculite, and perlite in a 2:1:1 ratio (v/v/v). The pH of the soil mixture was adjusted to 5.7-6.5 with dolomitic lime. The prepared soil mixture was incubated at room temperature for 48 hours, followed by baking at 70 °C for 48 hours. After germination, each cell was thinned to contain a single plant. Plants were fertilized with 1x Hoagland’s solution according to the following schedule: 2 mL/cell in the first week, 14 mL/cell in the second week, 28 mL in the third week, 44 mL in the fourth week. Plants were grown in a growth chamber under a 18 h light/6 h dark photoperiod with 180 µM m^−2^s^−1^ PPFD light provided by t5 fluorescent lamps with day/night temperatures of 24/18 °C and 60-65 % relative humidity.

### Touch treatments and thigmomorphogenesis analysis on soil

For touch treatments using a soft paintbrush, 10-day-old wild-type and mutant seedlings were gently brushed 10 times twice daily with a soft paintbrush at approximately 7-hour intervals. The number of brushings was later increased to 15 times per session when the seedlings were 17 days old, continuing the treatment until each plant reached flowering. Plants were considered to flower when their first flower opened. Thigmomorphogenetic changes were measured at flowering, with specific measurements depending on the *Arabidopsis* genotype and growth conditions. Rosette area was determined using Easy Leaf Area (Easlon and Bloom, 2014). LeafJ, an ImageJ plugin (Maloof et al., 2013), was used to measure petiole length, blade area, blade length, and blade width. Total rosette leaf area was measured using ImageJ (version: 1.53q). The percentage of flowering plants on a given day after planting was calculated by dividing the number of flowering plants on that day by the total number of plants.

For automated touch treatments, a fully automated machine, custom-built in the laboratory and equipped with a linear paintbrush bristle beam, was utilized for mechanical touch treatments. This regimen was initiated at 9:30 AM and consisted of three rounds per day, 8 hours apart. Within each 1-hour round, individual plants were touched 40 times by the bristles. The apparatus included a programmable controller/motor and a linear actuator, with the frame height adjusted according to plant height. Photographs of representative plants displayed in the figures were taken at flowering, when plants were considered ready for harvest or final assessment.

For touch treatments with MeJA, 11-day-old wild-type and *rrtf1* mutant seedlings were subjected to a single daily application of 50 μM MeJA or water as a control. Following application, plants were transferred to a separate growth chamber for up to 3 hours to allow for the dissipation of volatilized MeJA, thereby minimizing potential carryover effects.

In experiments involving a single touch event, 10- to 12-day-old seedlings were manually brushed back and forth four times with a soft paintbrush. Plant shoots were subsequently harvested at designated times for transcriptional and hormone analyses.

### RNA extraction, cDNA synthesis, and RT-qPCR

RNA was extracted from *Arabidopsis* shoot samples immediately using TRIzol (Invitrogen) or RiboZol (VWR) according to the manufacturer’s directions. Otherwise, samples were collected on ice and immediately frozen in liquid N_2_. Frozen tissue samples were ground into a fine powder using a mortar and pestle. RNA samples were digested with DNase I (NEB) and purified using Monarch RNA Cleanup Kit (NEB). cDNA was synthesized using ProtoScript II Reverse Transcriptase (NEB), followed by RNase H treatment (NEB). QPCR reactions were run using PowerUp Sybr Green Master mix (Applied Biosystems) on a Bio-Rad CFX96 Real-Time PCR Detection system. Gene expression was estimated using the 2^−ΔΔ*C*^_T_ method (Livak and Schmittgen, 2001) using 3 technical replicates for each of four biological replicates. *UBIQUITIN-CONJUGATING ENZYME 21* (*UBC21, AT5G25760*) was used for normalization. Primer sequences for the targeted genes are provided in Table S3 (Matsuo et al., 2015).

### RNA-seq library preparation, data processing, and differential gene expression analysis

Extracted RNA samples were treated with DNase I (NEB) and purified using Monarch RNA Cleanup Kit (NEB). RNA quality was evaluated by an Agilent 2100 Bioanalyzer. For RNA-seq analysis for temporal changes in Col-0, three biological replicates from each time point were used to construct sequencing libraries using the NEBNext® Ultra™ II Directional RNA Library Prep Kit for Illumina® (NEB). Libraries were examined by an Agilent 2100 Bioanalyzer prior to sequencing on an Illumina NovaSeq with 50 bp paired-end reads. For RNA-seq analysis comparing Col-0 and *rrtf1-1*, three biological replicates from each genotype/time point were used to construct sequencing libraries using the NEBNext® Ultra™ II RNA Library Prep Kit for Illumina® (NEB). Libraries were examined by an Agilent 2100 Bioanalyzer prior to sequencing on an Illumina NovaSeqX+ with 150 bp paired-end reads.

RNA-Seq data processing was conducted using the advanced computing resources at Texas A&M High Performance Research Computing. Approximately 10 to 57 million reads per sample were generated, trimmed using Trimmomatic (leading:3, trailing:3, slidingwindow:4:15, minlen: 36) to remove adapter sequences and low-quality reads, and aligned to the *Arabidopsis thaliana* reference genome (TAIR10) using HISAT2 (2.2.1-foss-2018b-Python-3.6.6) (Bolger et al., 2014; Kim et al., 2019). Mapping statistics are summarized in Table S1. The resulting files were sorted and filtered using SAMtools (1.10-GCC-9.3.0) (Li et al., 2009). Raw counts were generated by Subread (2.0.0-GCC-8.3.0) using featureCounts along with *Arabidopsis* genome annotation (Arabidopsis_thaliana.TAIR10.55.gtf) (Liao et al., 2013). The raw count data were analyzed using DESeq2 and edgeR to identify differentially expressed genes (DEGs, Robinson et al., 2010). For DESeq2-based analysis, DEGs were determined using the criteria: absolute value of log_2_ Fold Change > 1 and adjusted p-value (FDR) < 0.05, which was corrected using the R package ‘IHW’ (Love et al., 2014; Ignatiadis et al., 2016). For edgeR-based analysis, DEGs were determined using the criteria: absolute value of log_2_ Fold Change > 1 and false discovery rate (FDR) < 0.05 with the glmTreat function (Chen et al., 2025). Batch effects were removed using limma from raw counts of combined RNA-seq datasets (Ritchie et al., 2015).

### GO, TF, and TF binding site enrichment analysis

Gene Ontology (GO) analysis was performed using ShinyGO v0.82 and PANTHER (Protein ANalysis THrough Evolutionary Relationships) with an FDR cutoff of 0.05, a minimum pathway size of 10 and a maximum size of 2000 (Ge et al., 2020; Thomas et al., 2022). Transcription factor enrichment analysis was conducted via clusterProfiler (Wu et al., 2021). Fold enrichment was determined using the formula: GeneRatio (ratio of input genes annotated with a term) / BgRatio (ratio of all genes annotated with this term). TF2Network was used to search for transcription binding sites (Kulkarni et al., 2018).

### Promoter motif analysis

Promoter motif analysis was performed using MEME on genes upregulated at 10 min after touch, including 669 genes common to both Col-0 and the *rrtf1* mutant, 96 genes unique to Col-0, and 67 genes unique to the *rrtf1* mutant (Bailey and Elkan, 1994). One thousand base pairs of sequence upstream of the start ATG were analyzed in simple enrichment analysis of motifs (SEA) using a model of control sequences against the sets of unique genes (Bailey and Grant, 2021).

### Phytohormone analysis

For phytohormone analysis, three to four replicates of each sample (composed of 5-6 shoot sections) were collected on ice, weighed, and immediately frozen in liquid N_2_. Stable isotope-labeled standards for each hormone species (GA_1_, GA_4_, GA_5_, GA_8_, GA_20_, IAA, ABA, and JA) were added to the tissue samples, which were then extracted and semi-purified as described previously (Yao and Finlayson, 2015). Samples were subsequently derivatized with bromocholine (Kojima et al., 2009) and quantified by LC-MS/MS (Waters Aquity UPLC, Waters TQD) in positive MRM mode using the bromocholine derivative precursor/product transitions presented in Table S2.

### Exogenous MeJA and Jarin-1 application

To examine the effect of MeJA on *RRTF1* gene expression in the absence of mechanical stimulation, ten-day-old wild-type *Arabidopsis* seedlings grown on soil were exposed to MeJA. A 10 mM MeJA solution was dispensed into several open 1.5-mL tubes that were placed intermittently on the soil surface among the seedlings inside a sealed chamber, without direct plant contact. Plant shoots were harvested 15 minutes after the start of MeJA exposure. For RT-qPCR analysis, *Arabidopsis* seeds were surface sterilized with ethanol and bleach. The seeds were planted on plates containing 0.4X MS, 0.2 % sucrose, MES 0.5 g/L, bactoagar 8 g/L, and pH adjusted to 5.7. The seeds were planted on the media surface to have plants grow erect to avoid any contact impacts on touch signaling. The plates were stratified at 4 °C for 3 days before transferring to the growth chamber. Seven-day-old wild-type and mutant seedlings were applied by spraying 322.5 μL of water and 50 μM Jarin-1 on the plates. Plants were harvested 1 hour after application.

### Enrichment tests

Genome-wide binding profiles for RRTF1, WRKY18, and WRKY40 were identified using publicly available DAP-seq data from the *Arabidopsis* Cistrome Database (O’Malley et al., 2016; https://neomorph.salk.edu). To identify potential regulatory modules, the intersection between these transcription factor targets and the genes differentially expressed in response to mechanical stimulation was analyzed. The statistical significance of the overlap between gene sets was determined using a hypergeometric test (or Fisher’s exact test), with a significance threshold of *p* < 0.05.

### Statistical analysis

All statistical analyses were performed in R (version 4.2.2). Normality and homogeneity of variance were assessed using Shapiro-Wilk and Levene’s tests, respectively. Temporal variations in morphology, gene expression, and hormone abundances were analyzed using either two-way analysis of variance (ANOVA). However, given the dynamic range of the touch response, some datasets violated ANOVA assumptions; in these instances, generalized linear mixed models were used. Significant effects were further analyzed using Tukey’s HSD post-hoc tests. For pairwise comparisons between two specific groups, two-tailed Student’s t-tests were applied. Principal component analysis (PCA) of transformed read counts was performed using the plotPCA function in the DESeq2 R package.

## Data availability

The raw RNA sequencing data generated during this study are available in the NCBI Sequence Read Archive (SRA) under BioProject PRJNA1428196. All other data supporting the findings of this study are included in the paper and/or its Supporting Information. Custom scripts used for data analysis and figure generation are available at https://github.com/psk413/RRTF1-dependent_touch_transcriptome.git.

## Acknowledgements

This work is dedicated to the memory of the late Dr. Scott A. Finlayson, with profound gratitude for his mentorship and unwavering support. This work was supported by Texas A&M AgriLife Research (S.A.F.). S.P. was supported in part by a Tom Slick Fellowship from the College of Agriculture and Life Sciences, Texas A&M University. S.P. gratefully acknowledges Texas A&M AgriLife Research and the Department of Soil and Crop Sciences, Texas A&M University for their continued support following my graduation. Sequencing was performed at Texas A&M AgriLife Research and Novogene. Portions of this research were conducted with the advanced computing resources provided by Texas A&M High Performance Research Computing.

